# Discovery of anti-amoebic inhibitors from screening the MMV Pandemic Response Box on *Balamuthia mandrillaris, Naegleria fowleri* and *Acanthamoeba castellanii*

**DOI:** 10.1101/2020.05.14.096776

**Authors:** Christopher A. Rice, Emma V. Troth, A. Cassiopeia Russell, Dennis E. Kyle

## Abstract

Pathogenic free-living amoebae, *Balamuthia mandrillaris, Naegleria fowleri* and several *Acanthamoeba* species are the etiological agents of severe brain diseases, with case mortality rates >90%. A number of constraints including misdiagnosis and partially effective treatments lead to these high fatality rates. The unmet medical need is for rapidly acting, highly potent new drugs to reduce these alarming mortality rates. Herein, we report the discovery of new drugs as potential anti-amoebic agents. We used the CellTiter-Glo 2.0 high-throughput screening methods to screen the Medicines for Malaria Ventures (MMV) Pandemic Response Box in a search for new active chemical scaffolds. Initially we screened the library as a single-point assay at 10 and 1 µM. From these data, we reconfirmed hits by conducting quantitative dose response assays and identified 12 hits against *B. mandrillaris*, 29 against *N. fowleri* and 14 against *A. castellanii* ranging from nanomolar to low micromolar potency. We further describe 11 novel molecules with activity against *B. mandrillaris*, 22 against *N. fowleri* and 9 against *A. castellanii.* These structures serve as a starting point for medicinal chemistry studies and demonstrate the utility of phenotypic screening for drug discovery to treat diseases caused by free-living amoebae.

## 1. Introduction

Pathogenic free-living amoebae are highly lethal organisms whose under-recognized infections pose a significant risk to human health. *Balamuthia mandrillaris, Naegleria fowleri* and *Acanthamoeba* species are causative agents of encephalitis in humans as well as a variety of other species including, but not limited to, baboons, monkeys, dogs, mice, and bovines [1,2]. In spite of treatment, the fatality rate for human encephalitic disease caused by free-living amoebae remains >90% [3,4]. In addition, *Acanthamoeba* species can manifest as a cutaneous or keratitis infection; *Balamuthia mandrillaris* is also capable of manifesting as a cutaneous infection [1]. Drug discovery efforts against these amoebae have been scarce, though the encephalitic syndromes caused by them result in death the majority of the time. Further effort is warranted to find novel therapeutics.

*Balamuthia mandrillaris* was initially isolated by Visvesvara *et al.*, [5] in 1986 from the brain of a pregnant mandrill baboon. This protozoan parasite is presumed to occupy soil and freshwater environments and is capable of transforming into a highly-resistant ternate cyst under adverse conditions [2,6-8]. Nonetheless, contact with soil is considered a predisposing factor for these infections as *Balamuthia* has only been isolated from this source, making it a likely reservoir for this amoeba [8-10]. *B. mandrillaris* is an etiological agent for subacute or chronic *Balamuthia* Amoebic Encephalitis (BAE), the central nervous system (CNS) infection, as well as cutaneous and systemic infections in both animals and humans [2,11,12]. A key differentiating factor for BAE is that cases have not only been reported in immunocompromised hosts, but also in immunocompetent individuals with a higher frequency in young children and the elderly [12-17]. It is speculated that the route of entry for this parasite is either via a skin ulceration or the lower respiratory tract followed by hematogenous dissemination to the brain where the amoeba likely enters the CNS through the middle cerebral arterial supply to the choroid plexus [6,12,18].

The case fatality rate for BAE is ∼92%, with death resulting within one week to several months after the initial onset of symptoms [15,19]. Clinical manifestations of BAE include drowsiness, change in mental status or behavior, low-grade fever, headache, stiff neck, hemiparesis, aphasia, cranial nerve palsies, and seizures [6,13,14]. The US Centers for Disease Control and Prevention (CDC) recommends a multi-drug regimen that is based on a limited number of successful clinical cases and previously identified *in vitro* drug susceptibility. The therapeutic drug cocktail consists of pentamidine, sulfadiazine, flucytosine, fluconazole, azithromycin or clarithromycin, and recently, miltefosine, has been added [4,19]. With the poor success of clinical treatment and high mortality rates associated with disease, there is an urgent need for novel drug discovery for these deadly amoebae.

*Naegleria fowleri* is the only pathogenic species of the genus to humans. It is the causative agent of Primary Amebic Meningoencephalitis (PAM), a rapidly fatal infection. First identified as a pathogenic agent in 1965, *N. fowleri* has received minimal research attention though PAM cases are >97% fatal and highly publicized when they occur [11,20]. As a thermophilic free-living amoeba inhabiting both soil and freshwater, the number of individuals worldwide with the potential to be infected by *N. fowleri* is substantial. With global temperatures increasing, the habitable freshwater sources where *N. fowleri* can thrive have expanded as well. Typically, most PAM cases in the U.S. occur in the warmer southern-tiered states [1]. In the past 10 years, PAM cases have been reported in northern U.S. states, as far north as Minnesota, which previously had no reported cases [21]. PAM is characterized by non-specific symptoms such as headache, fever, nausea, and vomiting during the early stages of infection. Late stage symptoms include stiff neck, confusion, hallucinations, and seizures [22]. Death typically occurs 5-12 days after initial symptom onset [11]. With non-specific early symptoms and late stage symptoms that are identical to those of viral and bacterial meningitis, PAM is often misdiagnosed and thus likely under-reported.

Infection occurs when *N. fowleri*-contaminated water enters the nasal cavity. Most commonly, individuals are infected when swimming or playing in untreated freshwater, such as rivers and lakes, and improperly chlorinated swimming pools [23]. A significant number of cases have also linked to neti-pot use [21]. Upon entering the nasal cavity, amoebae that traverse the cribriform plate via the olfactory nerve find their way into the frontal lobe of the brain. In the brain, *N. fowleri* causes hemorrhage and incites a massive inflammatory response that ultimately results in patient coma and death [1,11]. A therapeutic combinational cocktail is the standard for PAM, the cocktail includes amphotericin B, an azole (ketoconazole, fluconazole, miconazole), azithromycin, rifampin, and more recently, miltefosine [4,23]. Induced hypothermia has been tried in addition to the therapeutic cocktail with variable success [21]. Little has changed in regards to PAM therapy since the 1970s. *N. fowleri* infection is still exceptionally lethal over 50 years since its initial discovery. Novel therapeutics for PAM must be developed to reduce the high mortality rate and prepare for the potential increase of PAM cases, as temperatures start increasing yearly.

In 1930, Castellani discovered *Acanthamoeba*, another genus of free-living amoebae found in freshwater and soil, to be potentially pathogenic [18]. *Acanthamoeba* species typically cause disease in immunocompromised patients but are also capable of facultatively infecting the cornea of immunocompetent contact lens wearers causing an extremely debilitating disease known as *Acanthamoeba* keratitis (AK). The symptoms of AK develop within a few days to weeks and consist of overactive tear production, photophobia, inflammation/redness, stromal infiltration, epithelial changes, oedema, stromal opacity, peripheral perineural infiltrates and incapacitating pain due to radial neuritis [24]. In immunocompromised individuals, the protist cannot only cause cutaneous amoebiasis and nasopharyngeal infections, but it can also disseminate to the CNS causing Granulomatous Amoebic Encephalitis (GAE) [25]. Similarly, to *B. mandrillaris, Acanthamoeba* is suspected to gain entry to the dead-end host via a skin lesion or the respiratory tract then through hematogenous dissemination to the brain [26]. Clinical manifestations of GAE include headache, fever, personality alterations, somnolence, hemiparesis, aphasia, diplopia, nausea, dizziness, cranial nerve palsies, seizures and coma [26,27]. The therapeutic measures utilized for infections caused by this parasite are laborious and ineffective, possibly due to the likelihood of inducing encystation and an inability to kill the double-walled resistant cyst stage that can be found in host tissues, unlike in PAM.

Though there are no drugs specifically approved for treatment by the FDA, the current treatment regimen for AK consists of a biguanide (polyhexamethylene biguanide [PHMB] or chlorhexidine) and a diamidine (propamidine or hexamidine) with hourly administration for 48 hours followed by a cessation of the nighttime dosing for 2 days, concluding with 3-4 weeks of treatment every 2 hours [24,28,29]. Even with this arduous treatment plan, there is the possibility for recrudescence of disease, toxic keratopathy, and vision impairment, with 2% of patients becoming blind [30]. The combinational chemotherapeutic regimen for GAE consists of an azole (ketoconazole, fluconazole, itraconazole or voriconazole), pentamidine isethionate, sulfadiazine, amphotericin B, azithromycin, rifampin, and miltefosine [4,11,31-33]. This cocktail of drugs remains ineffective even with the implementation of extreme measures including surgery and cryotherapy with most cases ending in death [1,31]. Furthermore, due to the renascent nature of the cyst stage, a longer period of treatment is needed which leads to additional trepidation for the development of drug resistance. Overall, the numbers of cases of AK have shown a significantly increasing trend worldwide in the past few decades with the estimated number of cases being up to 1.5 per 10,000 contact lens wearers depending on location [3, 34-39], thus confirming the need for continued efforts to discover more effective chemotherapeutics.

The Medicines for Malaria Venture (MMV) Pandemic Response Box is a drug library that was made available in January 2019 and is provided free of charge for research purpose. Thus far, 37 copies of this drug library have been distributed worldwide. The library consists of 400 compounds that are either already available on the market or are in various stages of drug discovery or development. It is comprised of 201 (50.25%) antibacterial compounds, 153 (38.25%) antiviral compounds, and 46 (11.5%) antifungal compounds. Previous high-throughput screening efforts on free-living amoebae have yielded promising leads for therapeutic development [40-47]. However, there is still a need for the discovery and development of novel chemical scaffolds potent against these amoebae. The diverse mechanisms of action (MOAs) of the compounds found within the Pandemic Response Box represent a promising opportunity for anti-amoebic drug discovery and a starting point for structure-based drug design (SBDD).

Herein we describe the trophocidal activity of 400 bioactive compounds from screening the MMV Pandemic Response Box independently against each of the pathogenic free-living amoebae: *B. mandrillaris, N. fowleri* and *A. castellanii.*

## 2. Results

### 2.1 Screening results for single point assays

A library of 400 drug-like compounds, with diverse MOAs, were assembled at 10 mM in dimethyl sulfoxide (DMSO) by Medicines for Malaria Venture (MMV, Switzerland) for open access screening against neglected and infectious diseases. Initially, each compound was screened at two concentrations, 10 and 1 µM, against pathogenic amoebae *B. mandrillaris* (Figure 1. A & B), *N. fowleri* (Figure 1. C & D) and *A. castellanii* (Figure 1. E & F). We used cell viability, determined with the CellTiter-Glo (CTG) 2.0 assay, as the endpoint for the single point screening assays for all amoebae. The single point assays yielded 43 compounds that were active (>33% inhibition) against *B. mandrillaris*, 104 compounds for *N. fowleri* and 24 compounds for A. castellanii. Of the 43 compounds identified against *B. mandrillaris*, 14 compounds (33%) were antifungals, 13 compounds (30%) were antivirals, and 16 compounds (37%) were antibacterials. For *N. fowleri* we identified activity of 24 antifungal compounds (23%), 32 antivirals (31%), and 48 antibacterials (46%). Of the 24 compounds identified against *A. castellanii*, 15 compounds (62%) were antifungals, 4 compounds (17%) were antivirals, and 5 compounds (21%) were antibacterials.

**Figure 1.**
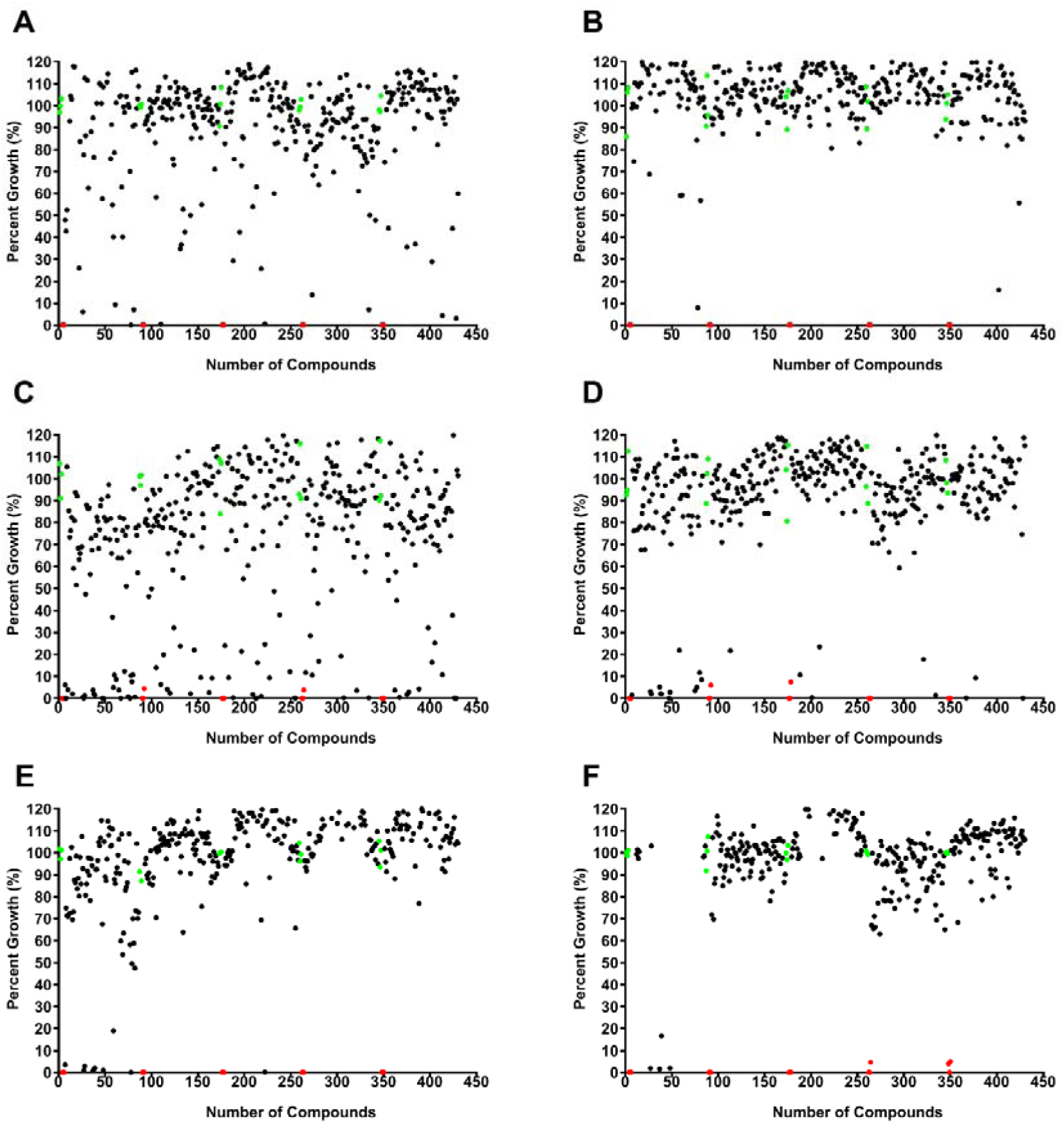
Single point screening assay of 400 compounds from the Medicines for Malaria Venture (MMV) Pandemic Response Box were carried out using pathogenic *B. mandrillaris* at 10 µM (**A**) and 1 µM (**B**) (n=1), *N. fowleri* at 10 µM (**C**) and 1 µM (**D**) (n=1), and *A. castellanii* at 10 µM (**E**) and 1 µM (**F**) (n=1). Each black circle represents an individual compound response, the red circles are positive controls (chlorhexidine, 12.5 µM), and the green circles are negative growth controls (0.1% and 0.01% DMSO for 10 µM and 1 µM plates, respectively) included on each plate.

### 2.2 Dose-response screening results

By using the CTG assay, we performed quantitative dose response assays to reconfirm the potency of hit compounds from single point screens against all amoebae. All hits that produced ≥33% inhibition in the primary screen were assessed with duplicate 2-fold serial dilutions to determine the 50% inhibitory concentration (IC_50_). We reconfirmed 12 hits against *B. mandrillaris* (Table 1), 29 against *N. fowleri* (Table 2) and 14 against *A. castellanii* (Table 3), with potencies ranging from nanomolar to low micromolar. We counter-screened all the reconfirmed hits for cytotoxicity 3-(4,5-dimethylthiazol-2-yl)-5-(3-carboxymethoxyphenyl)-2-(4-sulfophenyl)-2H-tetrazolium (MTS) assay against A549 lung carcinoma cells and identified only 10 compounds that displayed cytotoxicity from < 310 nM to 8 µM; all others tested were ≥10 µM. We determined the selectivity index (A549 IC_50_/Amoebae IC_50_), and defined an index of ≥10 to be our standard for further evaluation as a potentially useful drug for treatment of these parasitic diseases.

**Table 1.**
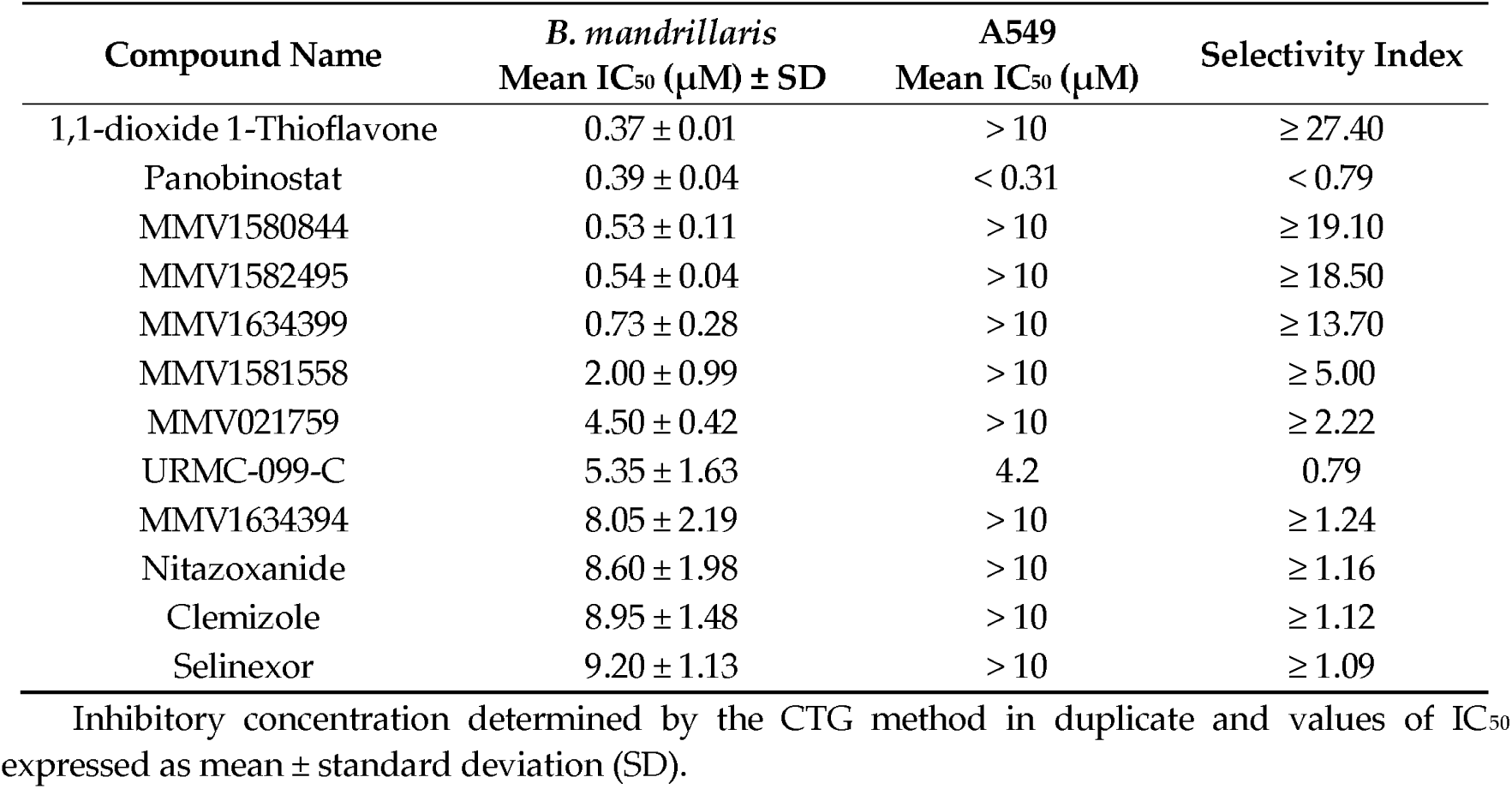
Dose response activity of confirmed hits against *B. mandrillaris*.

**Table 2.**
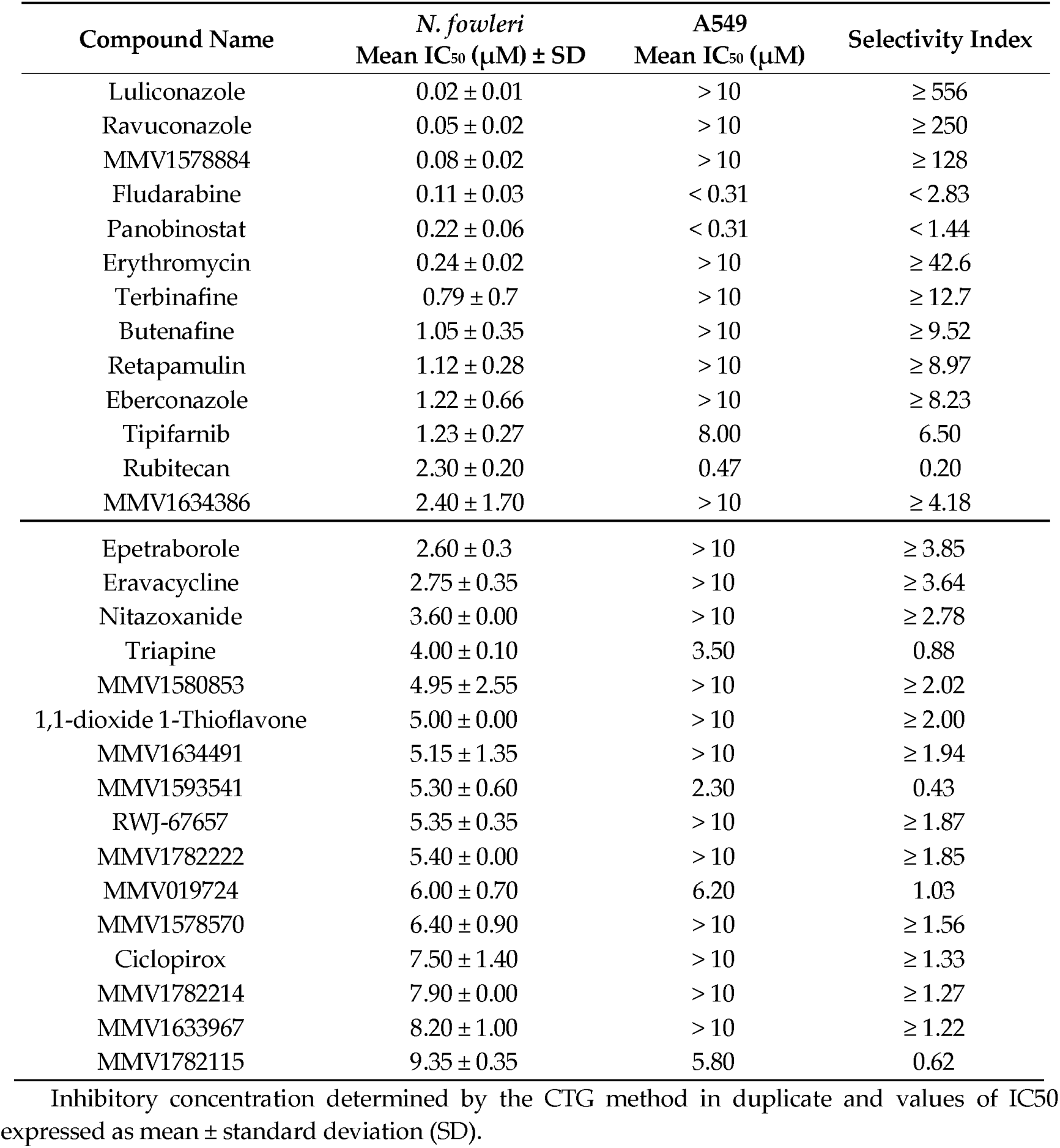
Dose response activity of confirmed hits against *N. fowleri*.

| Compound Name | *N. fowleri*<br>Mean $IC_{50}$ ($\mu\text{M}$) ± SD | A549<br>Mean $IC_{50}$ ($\mu\text{M}$) | Selectivity Index |
| --- | --- | --- | --- |
| Luliconazole | $0.02 \pm 0.01$ | $> 10$ | $\geq 556$ |
| Ravuconazole | $0.05 \pm 0.02$ | $> 10$ | $\geq 250$ |
| MMV1578884 | $0.08 \pm 0.02$ | $> 10$ | $\geq 128$ |
| Fludarabine | $0.11 \pm 0.03$ | $< 0.31$ | $< 2.83$ |
| Panobinostat | $0.22 \pm 0.06$ | $< 0.31$ | $< 1.44$ |
| Erythromycin | $0.24 \pm 0.02$ | $> 10$ | $\geq 42.6$ |
| Terbinafine | $0.79 \pm 0.7$ | $> 10$ | $\geq 12.7$ |
| Butenafine | $1.05 \pm 0.35$ | $> 10$ | $\geq 9.52$ |
| Retapamulin | $1.12 \pm 0.28$ | $> 10$ | $\geq 8.97$ |
| Eberconazole | $1.22 \pm 0.66$ | $> 10$ | $\geq 8.23$ |
| Tipifarnib | $1.23 \pm 0.27$ | 8.00 | 6.50 |
| Rubitecan | $2.30 \pm 0.20$ | 0.47 | 0.20 |
| MMV1634386 | $2.40 \pm 1.70$ | $> 10$ | $\geq 4.18$ |
| Epetraborole | 2.60 ± 0.3 | > 10 | ≥ 3.85 |
| Eravacycline | 2.75 ± 0.35 | > 10 | ≥ 3.64 |
| Nitazoxanide | 3.60 ± 0.00 | > 10 | ≥ 2.78 |
| Triapine | 4.00 ± 0.10 | 3.50 | 0.88 |
| MMV1580853 | 4.95 ± 2.55 | > 10 | ≥ 2.02 |
| 1,1-dioxide 1-Thioflavone | 5.00 ± 0.00 | > 10 | ≥ 2.00 |
| MMV1634491 | 5.15 ± 1.35 | > 10 | ≥ 1.94 |
| MMV1593541 | 5.30 ± 0.60 | 2.30 | 0.43 |
| RWJ-67657 | 5.35 ± 0.35 | > 10 | ≥ 1.87 |
| MMV1782222 | 5.40 ± 0.00 | > 10 | ≥ 1.85 |
| MMV019724 | 6.00 ± 0.70 | 6.20 | 1.03 |
| MMV1578570 | 6.40 ± 0.90 | > 10 | ≥ 1.56 |
| Ciclopirox | 7.50 ± 1.40 | > 10 | ≥ 1.33 |
| MMV1782214 | 7.90 ± 0.00 | > 10 | ≥ 1.27 |
| MMV1633967 | 8.20 ± 1.00 | > 10 | ≥ 1.22 |
| MMV1782115 | 9.35 ± 0.35 | 5.80 | 0.62 |

Inhibitory concentration determined by the CTG method in duplicate and values of IC<sub>50</sub> expressed as mean ± standard deviation (SD).

**Table 3.** Dose response activity of confirmed hits against *A. castellanii.*

| Compound Name | *A. castellanii* | A549 | Selectivity Index |
| --- | --- | --- | --- |
|  | Mean IC<sub>50</sub> (μM) ± SD | Mean IC<sub>50</sub> (μM) |  |
| Ravuconazole | 0.02 ± 0.01 | > 10 | ≥ 625 |
| Isavuconazonium | 0.09 ± 0.02 | > 10 | ≥ 105 |
| MMV1634386 | 0.16 ± 0.09 | > 10 | ≥ 62.50 |
| Terbinafine | 0.56 ± 0.28 | > 10 | ≥ 18 |
| MMV1582496 | 0.61 ± 0.35 | > 10 | ≥ 16.40 |
| Ketoconazole | 1.24 ± 0.80 | > 10 | ≥ 8.10 |
| Amorolfine | 1.63 ± 1.62 | > 10 | ≥ 6.13 |
| Trifluoroacetic acid | 1.97 ± 1.00 | > 10 | ≥ 5.10 |
| Butenafine | 1.97 ± 1.76 | > 10 | ≥ 5.08 |
| Alexidine | 2.40 ± 1.56 | 2.10 | 0.88 |
| MMV1634491 | 3.35 ± 0.07 | > 10 | ≥ 2.99 |
| Eberconazole | 3.70 ± 1.13 | > 10 | ≥ 2.70 |
| Furvina | 4.20 ± 0.85 | > 10 | ≥ 2.38 |
| MMV1782221 | 4.43 ± 1.00 | > 10 | ≥ 2.26 |

Inhibitory concentration determined by the CTG method in duplicate and values of IC<sub>50</sub> expressed as mean ± standard deviation (SD).

### 2.3 Comparison of compound activity against three pathogenic amoebae

We compared results for all of the reconfirmed hits against the three pathogenic amoebae to determine if there are any compounds with potential for treating multiple pathogenic amoeba disease indications. We identified three compounds with activity that overlapped between *B. mandrillaris* and *N. fowleri* (1,1-dioxide 1-Thioflavone, panobinostat and nitazoxanide) and six compounds overlapping between *N. fowleri* and *A. castellanii* (ravuconazole, terbinafine, butenafine, eberconazole, MMV1634386 and MMV1634491) (Figure 2.). We did not find *Balamuthia* and *Acanthamoeba* to share any active hits. No compounds produced >50% at 10 µM inhibition for all three pathogenic amoebae.

**Figure 2.**
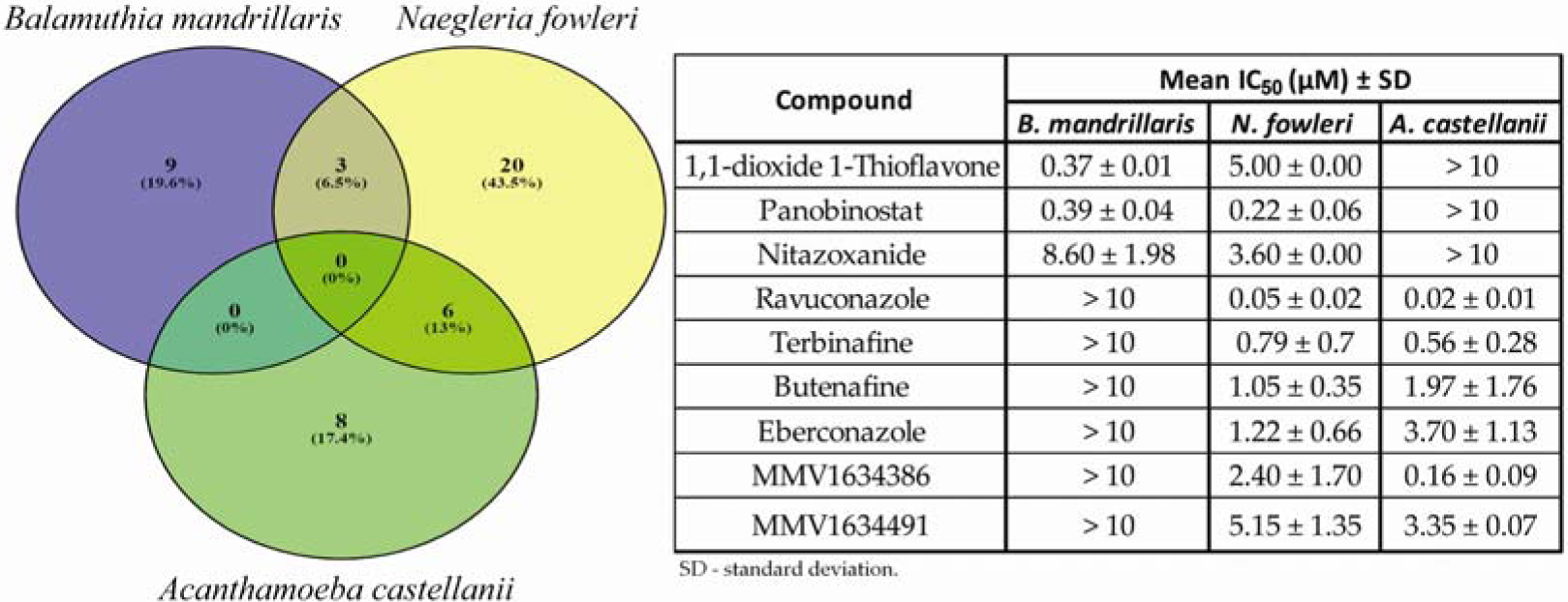
Venn diagram depicting the activity against *Balamuthia mandrillaris* (purple), *Naegleria fowleri* (yellow) and *Acanthamoeba castellanii* (green) of reconfirmed hits from the MMV Pandemic Response Box.

## 3. Discussion

There is a great unmet medical need for new therapeutics against the diseases caused by *B. mandrillaris, N. fowleri*, and *Acanthamoeba* spp. These are the epitome of neglected diseases with no major pharmaceutical companies and only a few academic labs working to discover new drugs. Phenotypic screening has significantly advanced the discovery and development of drugs targeting other parasitic protozoans and has been shown to be useful for discovery of new drugs against pathogenic free-living amoebae as well [42-44]. In this study we screened 400 bioactive compounds from the MMV Pandemic Response Box for activity against all three pathogenic free-living amoebae. Given the lack of drug discovery against these pathogens, the overarching goal is to find compounds with potential for treating multiple disease indications caused by these amoebae. In addition, we aimed to discover new drugs that could be repurposed as well as new starting points for drug discovery and lead optimization. We were successful in discovering 58 new leads from this rich resource of bioactive compounds, with some having activity against at least two of the amoebae pathogens.

After reproducibly reconfirming hits via quantitative dose response, we identified 12 compounds with IC_50_s <10 µM against *B. mandrillaris*. Compounds with nanomolar potency included 1,1-dioxide 1-Thioflavone, Panobinostat, MMV1580844, MMV1582495, and MMV1634399. Pharmacological properties for 1,1-dioxide 1-Thioflavone implicate its usage as an anticarcinogenic as well as an antimicrobial agent, with specific antiviral properties against cytomegalovirus and coxsackievirus [48]. Panobinostat, a histone deacetylase (HDAC) inhibitor was active against the amoeba, but it was not selective with toxicity at very low concentrations [49]. Though the molecule is approved and registered as a combinational chemotherapeutic to treat multiple myeloma, the mutagenicity and genotoxicity along with the poor pharmacokinetic profile of the drug significantly decreases the viability of the drug for amoebae, but suggests that HDAC might be a target for future studies [50]. MMV1581558, MMV021759, URMC-099-C, MMV1634394, nitazoxanide, clemizole, and selinexor all demonstrated IC_50_s ≤10 µM against *B. mandrillaris*. The mixed lineage kinase type 3 inhibitor, URMC-099-C, is known to induce amyloid-beta clearance in mouse Alzheimer’s models [51,52]. Though shown to be toxic, the brain-penetrating property of this molecule makes it an interesting lead compound for future drug discovery efforts. Nitazoxanide is FDA-approved and licensed to treat cryptosporidial diarrhea and other intestinal parasitic infections [53]. This thiazolide agent interferes with the electron transfer reaction in anaerobic metabolism and has been found to both inhibit Ebolavirus growth and increase the host antiviral response [54,55]. Griffin et al., [56] found that the benzimidazole histamine H1 antagonist, clemizole, acts as an antiepileptic and speculate that it could be used to treat Dravet syndrome based upon their study in zebrafish. Clemizole has also been implicated as a potential therapeutic against hepatitis C and shows anti-tumor as well as anti-allergic activities [57]. Selinexor acts by inhibiting the nuclear export protein, exportin 1, and has received FDA approval for the treatment of multiple myeloma in the U.S. [58].

In total, 29 compounds reproducibly yielded IC_50_ values ≤10 µM for *N. fowleri*. The majority (22 of 29) of the hit compounds have not been previously reported in the literature for activity against *N. fowleri*. The Pandemic Response Box screen yielded 7 active compounds with nanomolar potency against *N. fowleri*: luliconazole, ravuconazole, CRS-3123, fludarabine, panobinostat, erythromycin, and terbinafine. Aside from fludarabine and panobinostat, these compounds all generated selectivity indices ≥10, preliminarily indicating little cytotoxicity to A549 mammalian cells. Luliconazole and ravuconazole are novel compounds whose activity has not been previously described against *N. fowleri*. Luliconazole is an FDA-approved imidazole antifungal agent that is typically administered in a topical cream, and primarily prescribed for the fungal foot infection, tinea pedis [59]. Currently, luliconazole is being investigated for its use against a broad spectrum of fungal afflictions including aspergillosis, dermatophytosis, and onychomycosis [60]. Ravuconazole is a triazole antifungal approved for clinical use in Japan for treatment of tinea pedis but has yet to receive FDA-approval [61]. *In vivo* studies have demonstrated promising results for oral and intravenous bolus administration of ravuconazole. Groll *et al.*, [62] have shown ravuconazole is able to penetrate the blood-brain barrier (BBB) and reach brain tissue in an *In vivo* rabbit study. Ravuconazole has also been effective *in vivo* in models for disseminated candidiasis, intracranial cryptococcus, and invasive pulmonary aspergillosis [63-65]. Eberconazole was also identified as a hit compound against *N. fowleri*, making this the first report of its activity against this parasite. Eberconazole is an imidazole antifungal that is currently not FDA-approved. Eberconazole has shown promising in vitro results for dermatophytosis and candidiasis [66].

Luliconazole, ravuconazole, and eberconazole function by blocking ergosterol biosynthesis via inhibition of 14α-demethylase [67]. The azole class of drugs has long been known to be active against *N. fowleri* and an azole has been included in the treatment regimen since the 1980s [68]. Furthermore, 14 α-demethylase has been confirmed as a drug target against *N. fowleri* [69]. Posaconazole is the most recently described azole with promise as a therapeutic lead, and is implicated as an effective combinational partner in the treatment for PAM [42]. Both luliconazole and ravuconazole produced lower IC_50_ values than the reported posaconazole IC_50_, thus these compounds warrant further in vitro and *in vivo* evaluation.

CRS-3123, fludarabine, panobinostat, erythromycin, and terbinafine were also identified as nanomolar inhibitors of *N. fowleri*. We rediscovered these compounds that have been described previously in high-throughput screens [42,44]. CRS-3123 is an antibacterial compound that targets methionyl tRNA-synthetase and is currently involved in clinical trials for the treatment of *Clostridium difficile* [70]. Fludarabine is a purine nucleoside analog antineoplastic with FDA approval that is used in the treatment of leukemia and lymphoma [71]. Panobinostat is an FDA-approved histone deacetylase inhibitor antineoplastic used in the treatment of multiple myeloma [49]. Erythromycin is an FDA-approved antibacterial macrolide with a broad spectrum of clinical applications [72]. Terbinafine is an amine antifungal that inhibits squalene epoxidase and also is FDA-approved [73]. Though not nanomolar hits, butenafine and repatamulin also were reconfirmed as hit compounds ≤10 µM in this screen. Reconfirmation of hits identified from previous screening demonstrates the robustness of our assay and the effectiveness of our high-throughput screening methods.

For *A. castellanii*, 14 compounds yielded reproducible IC_50_s ≤10 µM. Of the hit compounds, 9 of 14 have not been previously described in the literature for activity against *Acanthamoeba* species. Our screen identified 5 inhibitors with nanomolar potency against *A. castellanii*: ravuconazole, isavuconazonium, MMV1634386, terbinafine, and MMV1582496. Additionally, all 5 compounds presented a selectivity index ≥10 and thus do not appear to display cytotoxic effects against A549 mammalian cells in vitro. Of these 5 compounds, the 3 with known MOAs target ergosterol biosynthesis, with ravuconazole and isavuconazonium specifically inhibiting CYP51, and terbinafine inhibiting squalene 2,3-epioxidase [74-76]. Ravuconazole, an inhibitor we discovered for *N. fowleri*, also is effective at inhibiting growth of a clinical isolate of fungal keratitis caused by *Scedosporium apiospermum*, and thus could have valuable potential in treating ocular infections caused by *Acanthamoeba* species [77]. Isavuconazonium, the pro-drug of isavuconazole, has also been shown to penetrate the BBB and is approved by the FDA to treat aspergillosis as well as mucormycosis [78-79]. The allylamine antifungal, terbinafine, has been used previously to treat a case of osteo-cutaneous acanthamoebiasis [80]. Compounds with IC_50_’s ≤10 µM against *A. castellanii* included ketoconazole, amorolfine, trifluoroacetic acid, butenafine, alexidine, MMV1634491, eberconazole, furvina, and MMV1782221. Ketoconazole, amorolfine, butenafine, and eberconazole interfere with the ergosterol biosynthesis pathway by inhibiting CYP51A1, Δ7,8-isomerase and the C14-reductase, squalene epoxidase, as well as 14α-demethylase, respectively [81-83]. Although not approved in the U.S. or Canada, amorolfine is approved and commonly used topically to treat dermatophyte infections (tinea capitis, tinea pedis, and onchomycosis) in Australia and the United Kingdom [84,85]. The benzylamine antifungal, butenafine, is approved by the FDA for topical treatment of tinea pedis and has also been found to be effective against *Leishmania* and a variety of ocular pathogenic fungal infections [86-88]. The antimicrobial activity of trifluoroacetic acid—commonly utilized as an antiseptic provided by the various ester groups found on the molecule [89]. Alexidine, being a bis-biguanide, acts via phase separation as well as the interruption of domain formation in membrane lipids and has been shown to have activity against several *Acanthamoeba* species [90]. The synthetic nitrovinylfuran broad spectrum antibiotic, furvina, was developed in Cuba and has shown antimicrobial activity against bacteria, fungi and yeasts by preferentially inhibiting protein synthesis at the P-site of the 30s ribosomal subunit [91]. We found ketoconazole, terbinafine, and alexidine to have reconfirmed its activity against *Acanthamoeba* as well as isavuconazonium and butenafine, both identified in a previous drug susceptibility screen [44,92-94]. These hits against *Acanthamoeba* further substantiate the robust nature of our high-throughput drug susceptibility screening techniques.

Unfortunately, no compounds had shared activity between all three amoebae at the final screening concentration tested of ≤10 µM. This could be due to our stringent activity criteria implemented to detect moderately active molecules that directly target the amoebae. Our screen would not identify immune modulators or compounds that target host processes that could affect amoeba infections. The results of this study as well as a large screen of 12,000 compounds suggest it is unlikely a potent compound (<1 uM) with pan-activity against the three-amoebae will be found without undergoing several rounds of structure-activity relationship (SAR) medicinal chemistry optimization. We did however identify three compounds that were shared between *B. mandrillaris* and *N. fowleri* (1,1-dioxide 1-Thioflavone, panobinostat and nitazoxanide) and six compounds that were shared between *N. fowleri* and *A. castellanii* (ravuconazole, terbinafine, butenafine, eberconazole, MMV1634386 and MMV1634491). Of these, ravuconazole, terbinafine, butenafine, eberconazole for *N. fowleri* and *A. castellanii* and panobinostat and nitazoxanide for *B. mandrillaris* and *N. fowleri* should be investigated further for *In vivo* efficacy and the potential off-label use for the various clinical diseases caused by these amoebae.

## 4 Materials and Methods

### 4.1. Maintenance of amoebae

#### 4.1.1. Balamuthia mandrillaris

Pathogenic *Balamuthia mandrillaris* (CDC:V039; ATCC 50209), a GAE isolate, isolated from a pregnant baboon at the San Diego Zoo in 1986 was donated by Luis Fernando Lares-Jiménez ITSON University, Mexico [43]. Trophozoites were routinely grown axenically in BMI media at 37°C, 5% CO_2_ in vented 75 cm^2^ tissue culture flasks (Olympus), until the cells were 80-90% confluent. For sub-culturing, 0.25% Trypsin-EDTA (Gibco) cell detachment reagent was used to detach the cells from the culture flasks. The cells were collected by centrifugation at 4,000 rpm at 4°C. Complete BMI media is produced by the addition of 10% fetal bovine serum (FBS) and 125 µg of penicillin/streptomycin antibiotics.

#### 4.1.2. Naegleria fowleri

Pathogenic *Naegleria fowleri* (ATCC 30215), a clinical isolate obtained from a 9-year old boy in Adelaide, Australia, that died of PAM in 1969 was previously purchased from the American Type Culture Collection (ATCC) [40]. Trophozoites were routinely grown axenically at 34°C in Nelson’s complete medium (NCM) in non-vented 75 cm^2^ tissue culture flasks (Olympus), until the cells were 80-90% confluent. For sub-culturing, cells were placed on ice to detach the cells from the culture flasks. The cells were collected by centrifugation at 4,000 rpm at 4°C. Complete NCM media is produced by the addition of 10% FBS and 125 µg of penicillin/streptomycin antibiotics.

#### 4.1.3. Acanthamoeba castellanii

Pathogenic *Acanthamoeba* castellanii T4 isolate (ATCC 50370) used in these studies was isolated from the eye of a patient in New York, NY, in 1978. This isolate was also purchased from ATCC. Trophozoites were routinely grown axenically at 27°C in Protease Peptone-Glucose Media (PG) in non-vented 75 cm^2^ tissue culture flasks (Olympus), until the cells were 80-90% confluent. For sub-culturing, cells were mechanically harvested to detach the cells from the culture flasks [95]. The cells were collected by centrifugation at 4,000 rpm at 4°C. Complete PG media is produced by the addition of 125 µg of penicillin/streptomycin antibiotics.

All experiments were performed using logarithmic phase trophozoites.

### 4.2. Compound library

The Pandemic Response Box (https://www.mmv.org/mmv-open/pandemic-response-box) was modelled after the previously successful Malaria and Pathogen Boxes, and is an open-source drug library comprised of 400 diverse compounds that includes: 201 antibacterial inhibitors (50.25%), 153 antiviral inhibitors (38.25%) and 46 antifungal inhibitors (11.5%). Compounds within this collection are either FDA approved and currently available on the pharmaceutical market or in earlier different stages of drug development. All stock compounds were supplied as 10 mM in DMSO.

### 4.3. In vitro CellTiter-glo trophocidal assay

The trophocidal activity of compounds was assessed using the CellTiter-Glo 2.0 luminescent viability assay (Promega, Madison, WI), as previously described [40,43]. Trophozoites were routinely cultured as described above and only logarithmic trophozoites were used. In brief, *B. mandrillaris, N. fowleri* or *A. castellanii*, trophozoites cultured in their corresponding media were seeded at 16,000, 3,000 or 1,440 cells/well into white 96-well plates (Costar 3370), respectively. Initially all compounds were assessed in a single point drug screen at 10 and 1 µM, as previously described. Inhibitors were assessed for percent inhibition using the criteria of ≤33% growth inhibition (no inhibition), 33-67% growth inhibition (moderate inhibition) and ≥67% growth inhibition (strong inhibition). Control wells were supplemented with 0.1% (10 µM screening plate) or 0.01% (1 µM screening plate) DMSO, as the negative controls, or 12.5 µM of chlorhexidine, as the positive control. All inhibitors that met the predetermined criteria of inhibiting parasite growth by ≥33% were next confirmed in quantitative dose response assays. Drugs were cherry-picked, dissolved in the media specific to each parasite, and assessed in 2-fold serial dilutions from the highest concentration of 10 µM. All assays were incubated at each of the parasites’ representative growth temperatures, described above, for 72 hours. At the 72-hour time point, 25 µl of CellTiter-Glo 2.0 reagent was added to all wells of the 96-well plates using the Biomek NX^P^ automated workstation. The plates were protected from light and contents were mixed using an orbital shaker at 300 rpm at room temperature for 2 minutes to induce cell lysis. After shaking, the plates were equilibrated at room temperature for 10 minutes to stabilize the luminescent signal. The ATP luminescent signal (relative light units; RLUs) were measured at 490 nm with a SpectraMax I3X plate reader (Molecular Devices, Sunnyvale, CA). Drug inhibitory concentration (IC_50_) curves were generated using total ATP RLUs where controls were calculated as the average of replicates using the Levenberg-Marquardt algorithm, using DMSO as the normalization control, as defined in CDD Vault (Burlingame, CA, USA). Values reported are from a minimum of two biological replicates with standard deviations.

### 4.4. Cytotoxicity screening of reconfirmed hits

Cytotoxicity of the reconfirmed hits was determined by using the CellTiter 96^®^ AQ_ueous_ One Solution Cell Proliferation Assay (Promega, Madison, WI) on A549 human lung carcinoma cells [40]. A549 cells were seeded at a concentration of 1.6 × 10^4^ cells/ml in 96-well tissue culture plates (Corning, NY), in the presence of serially diluted active hits against *B. mandrillaris, N. fowleri* or *A. castellanii*. Positive control wells contained cells and media; negative control wells contained 0.1% DMSO. Cells were grown in F12K medium supplemented with 10% FBS and 1% gentamycin (all supplied from Fisher Scientific, Atlanta, GA). The inhibitor concentration started at 10 µM and was diluted in doubling dilutions to assess cytotoxicity in comparison to the respective free-living amoeba. Total volume of each well was 100 µl and plates were incubated at 37°C, 5% CO_2_ for 72 hours. 4 hours before the time-point, 20 µl of MTS (Promega, Madison, WI) was added to each well. Inhibition of A549 growth was assessed at the 72-hour time-point measuring the optical density (OD) values at 490 nm using a SpectraMax I3X plate reader (Molecular Devices, Sunnyvale, CA). Curve fitting using non-linear regression was carried out using the average of replicates and the Levenberg-Marquardt algorithm, using DMSO as the normalization control, as defined in CDD Vault (Burlingame, CA, USA).

From these data we calculate a selectivity index (SI), SI = (IC_50_ A549)/(IC_50_ Amoeba). An SI value ≥10 is considered the standard for further evaluation as a potentially useful drug.

### 4.5. Statistical analysis

We used the Z’ factor as a statistical measurement to assess the robustness of our high-throughput screening assays (Figure 1) [96]. This factor uses the mean and standard deviation values of the positive and negative controls to assess data quality. The robustness of all of the plates screened had an excellent Z’-score value of 0.8 or above.

## 5. Conclusions

In conclusion, we identified several new chemical species with nanomolar to low micromolar potency against *B. mandrillaris, N. fowleri* or *A. castellanii*, which can be used as repurposed drugs or as starting scaffolds for medicinal chemistry to develop better therapeutics against these pathogenic free-living amoebae.

## Author Contributions

Conceptualization, C.A.R. and D.E.K.; methodology, C.A.R., E.V.T., and A.C.R.; validation, C.A.R., E.V.T., and A.C.R.; formal analysis, C.A.R., E.V.T., and A.C.R.; resources, D.E.K.; data curation, C.A.R., E.V.T., and A.C.R.; writing—original draft preparation, C.A.R., E.V.T., and A.C.R.; writing—review and editing C.A.R., E.V.T., A.C.R., and D.E.K.; visualization, C.A.R., E.V.T., and A.C.R.; supervision, C.A.R. and D.E.K.; funding acquisition, D.E.K. All authors have read and agreed to the published version of the manuscript.

## Acknowledgments

We would like to thank Medicines for Malaria Venture (MMV) for the curation and distribution of this drug library (MMV Pandemic Response Box).

